# Physiological and Biochemical Analyses of *Sorghum* Varieties Reveal Differential Responses to Salinity Stress

**DOI:** 10.1101/720789

**Authors:** W Mbinda, M Kimtai

**Affiliations:** Department of Biochemistry and Biotechnology, Pwani University, Kilifi, Kenya

**Keywords:** Chlorophyll content, Salinity stress, Salt tolerance, *Sorghum*, Relative water content

## Abstract

Salinity is among the most severe and widespread environmental constrains to global crop production, especially in arid and semi-arid climates and negatively affecting productivity of salt sensitive crop species. Breeding and selection of salt tolerant crop varieties is therefore necessary for sustainable plant productivity. Given that germination and seeding phases are the most critical phase in the plant life cycle, this study aimed to evaluate seed germination potential and associated traits under salt stress conditions as a simple approach to identify salt tolerant *Sorghum* varieties. There *Sorghum* varieties whose adaptation to various agroclimatic conditions is not well elucidated. Salinity stress was applied by addition of NaCl at three different levels of stress (100, 200 and 300 mM NaCl), while plants irrigated with water were used as controls. Evaluation of tolerance was performed on the basis of germination percentage, shoot and seed water absorbance, shoot and root length, leave water content, seedling total chlorophyll content and morphologic abnormality. Our results showed that salinity stress significantly impacts all features associated with germination and early development of seedlings. Our results indicated that that salinity stress substantially affects all traits associated with germination and early seedling growth, with the effect of salinity being dependent on the variety used and level of salinity stress applied. Among the tested *Sorghum* varieties, Gadam was established to the most salt tolerant variety, suggesting its potential use for cultivation under salinity stress conditions as well as its suitability for use as germplasm material in future *Sorghum* breeding programmes. For a greater insight into comprehensive mechanisms of salinity tolerance in *Sorghum*, we suggest further research on genomic and molecular analysis.

## INTRODUCTION

Salinity stress is a major constrain that affect crop growth and metabolism, resulting to severe damage and a loss of productivity primarily in arid and semi-arid regions [1]. Exposure to salinity stress triggers a variety of biochemical and physiological responses in plants and these responses include chlorophyll degradation, reduction in water content as well as morphological changes [2].

Germination is one of the most critical periods for a crop subjected to salinity. Higher salt stress retards seed germination and root emergence and leads to poor crop establishment which is deleterious and prevents the plant in maintaining their proper nutritional requirements necessary for their healthy growth [3]. Reduced germination is the consequence of either direct toxic effect of salt or the general delay of the germination process caused by osmotic stress [4].

*Sorghum* is a model crop for a more concerned crop improvement program in agriculture to utilize marginal lands, to meet energy and food demands which might be increased in the near future [5]. *Sorghum* is considered as a model crop for crop improvement for utilization of marginal lands in order to meet food demands which is expected to increase in the near future due to growing world population [6,7]. Even through *Sorghum* is mainly grown in poorly irrigated and partly saline conditions throughout the world, it is exposed to great deal of salt stress. Although much supporting evidences on biochemical and physiological responses under salinity are available, there is no specific information pertinent to *Sorghum* varieties grown in Kenya.

## MATERIALS AND METHODS

### Plant growth and stress treatment

Seeds of three finger millet varieties (Serena, Seredo and Gadam) provided by the Kenya Seed Company, Nairobi, Kenya were used as plant material. These varieties’ responses to salinity stress at germination and seedling growth period are have not been established. The healthy seeds were sorted by handpicking before washing them with distilled water to remove dust and other particles. Seeds were sown to a depth of approximately 1 cm in plastic pots that had been filled with sterile soil sand and perforated at the bottom for drainage. The pots were irrigated with different concentrations of NaCl (100, 200 and 300 mM NaCl) at an interval of 3 days for two weeks. Control seeds were irrigated with distilled water. Observations on the rate of germination were recorded on the 17th day of treatment. Seeds were considered to have germinated when the radicle was at approximately 2 mm long. The experiment was repeated three times with five replications for each treatment.

### Growth conditions upon salinity treatment

Germinated *Sorghum* seeds were grown for two weeks under greenhouse conditions. To find out the water deficit effects on growth of *Sorghum* seedlings, the seedlings were subjected to osmotic stress by irrigating with different concentrations NaCl (100, 200 and 300 mM NaCl) for 21 days at an interval of 3 days while the control pots were watered with distilled water. In each experiment, three replications were used for each set of treatment. After treatment, three plants from each treatment were sampled at random and the growth of the plants studied by recording the shoot length and root length.

### Relative water content estimation

One leaflet from the first fully expanded leaf of three plants per variety and per treatment was cut from a plant on the 21st day. Immediately after cutting, the leaflet was weighed to obtain the fresh weight (FW). Thereafter, the leaflet was immersed in double distilled water and incubated under normal room temperature for 4 hours. Afterwards, the leaflet was taken out, thoroughly wiped to remove the water on the blade surface and its weight measured to obtain turgid weight (TW). The leaflet was dried in an oven for 48 hours and its dry weight (DW) measured. The relative water content (RWC%) was calculated using the formula: RWC=[(FW−DW)/(TW−DW)] × 100.

### Determination of total chlorophyll content

To determine the chlorophyll amount, fresh leaves (0.2 g) of leaves plants were crushed in 80% acetone. Grinding was done by vortexing several times to remove chlorophyll efficiently. The extract was centrifuged at 5000 g for 3 minutes. The absorbance of the obtained supernatants was measured at 646 and 663 nm.

The total chlorophyll content in each sample, expressed in mg g−1 (FM), was calculated using the following formula: TC=20.2(A646)+8.02(A663) × V/1000 × W where V corresponds to the volume of total extract per liter and W is the mass of the fresh material [8].

### Statistical analysis

Data were analysed with Minitab statistical software version 19. One-way ANOVA tested for the significance of the salinity effects of germination percentage, plant growth characteristics, relative water content and total chlorophyll content. Differences between means were compared using the Fisher’s least significant difference test. Differences were considered significant when p<0.05.

## RESULTS

### Effects of salt stress on seed germination

The effect of NaCl stress on *Sorghum* seeds germination, evaluated by the percentage of germinated seeds after 17 days, is as shown in (Figure 1). The results show that for all *Sorghum* varieties, the germination rate decreased with an increase of the NaCl concentration. However, the negative effect of NaCl stress differed according to the varieties.

**Figure 1:**
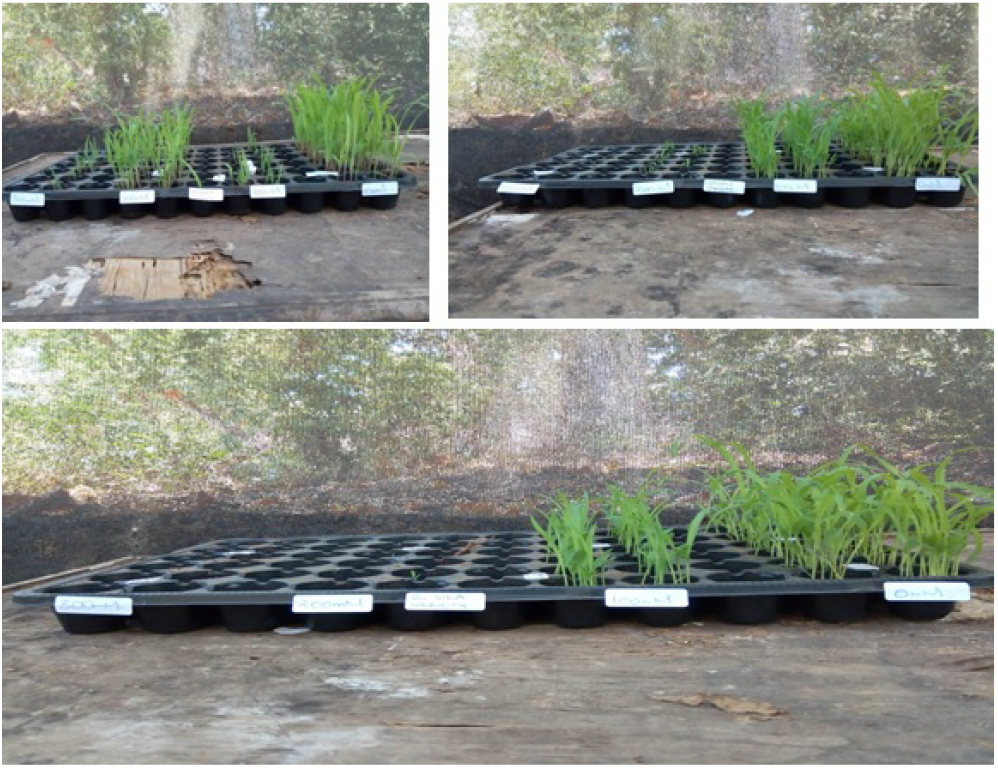
Effects of salinity stress upon germination of *Sorghum* varieties. (A) Serena (B) Gadam and (C) Sc Sila.

Under untreated conditions, results showed all the varieties had statistically similar germination rates ranging from 74.68% for Gadam to 75.13% for Serena (Figure 2). On 100 mM Gadam recorded a higher germination rate of 45 46% as compared to Sc Sila (43.61%) and Serana (34.62%.) At severe osmotic pressure of 300 mM NaCl, only 14.31% germination rate was recorded from Gadam, while Serena and Sc Sila recorded 6.93% and 2.66% respectively (Figure 2).

**Figure 2:**
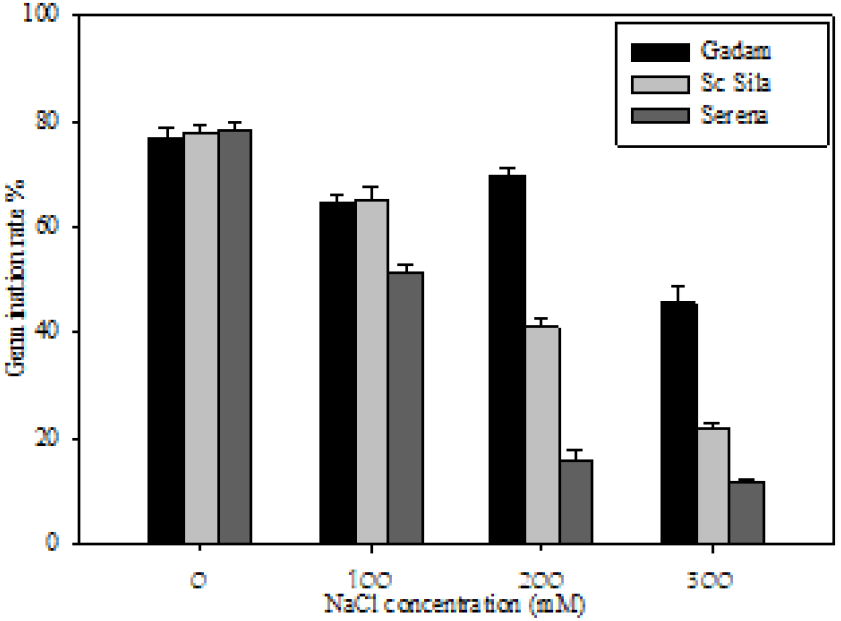
Effects on salinity stress on germination of *Sorghum* seeds.

### Effect of NaCl on seedlings growth

Analysis revealed statistically significant differences in root and shoot length among varieties and treatments (Figure 3). Although the increased concentrations of NaCl resulted in a significant reduction of both root and shoot length, these tissue types responded differently under salinity stress (Tables 1 and 2). Specifically, there was a reduction in length at NaCl concentrations of 100 mM for roots and shoots, respectively. However, root length did not provide an accurate estimate for the classification of varieties with respect to tolerance as the length across the varieties under similar NaCl treatments were similar. In relation to shoot length, Gadam was the best performing variety, whereas Serena and Sc Sila had poor growth of shoots when grown with 300 mM NaCl (Table 1). At control conditions, the *Sorghum* varieties had similar height. Similar results were also reported for root development under control and treatment conditions (Table 2). Despite the drastic effects on seedling elongation, no morphological deformities were observed, indicating that salinity stress does not result to the development of abnormal phenotypes.

**Figure 3:**
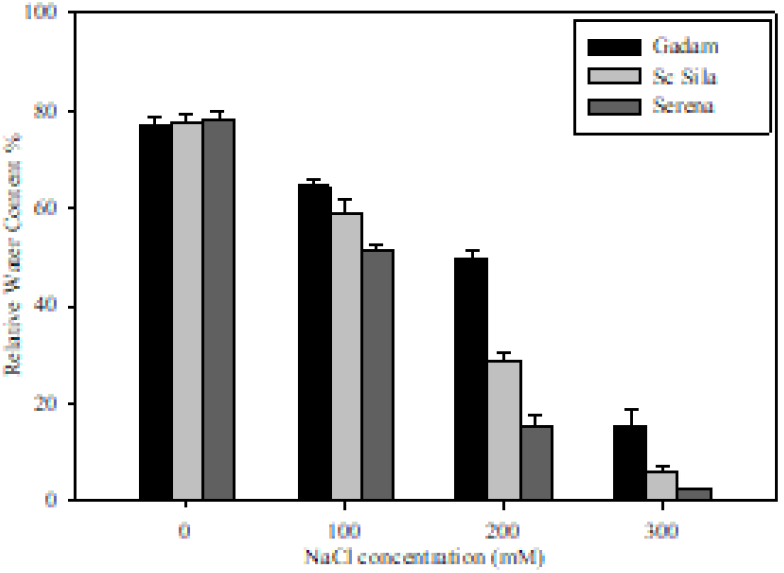
Effect of salt stress on leaf relative water content.

**Table 1:**
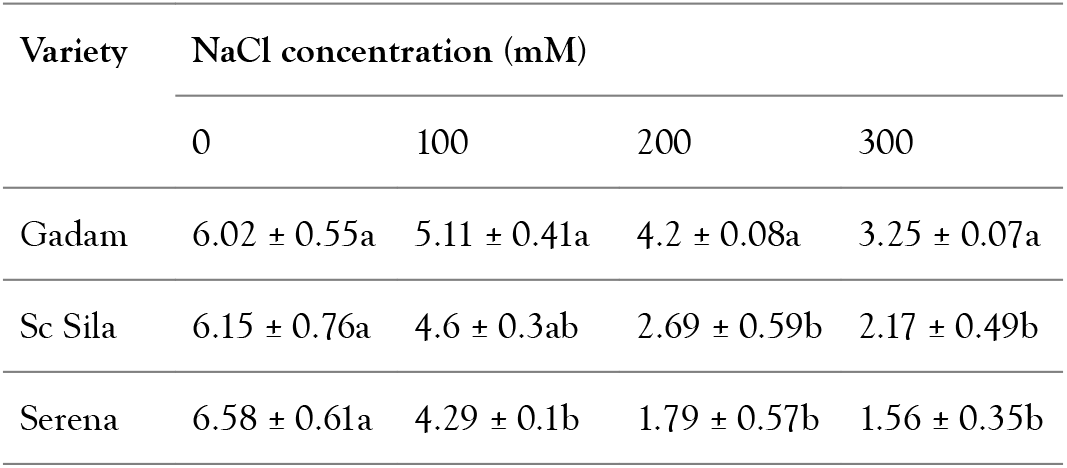
Effect of NaCl concentrations on shoot height and root length of different varieties of *Sorghum* seedlings.

**Table 2:**
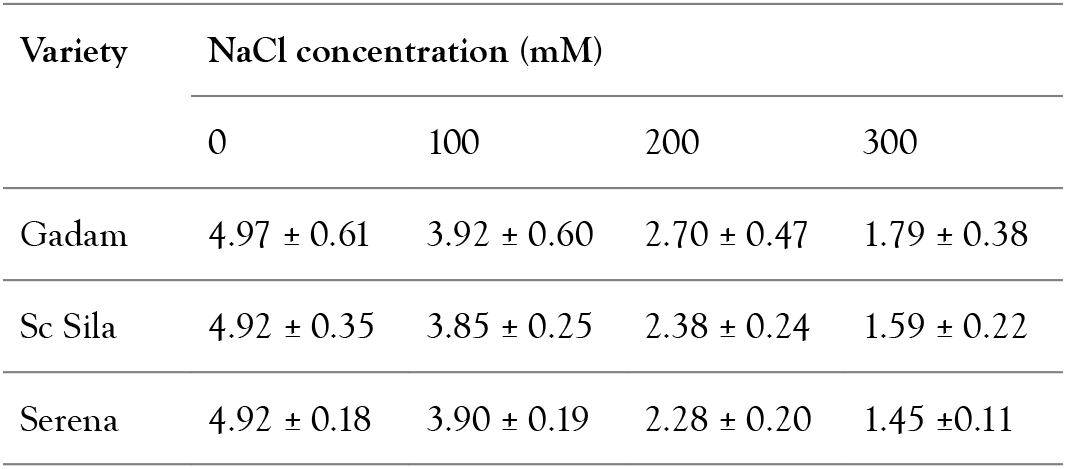
Effect of NaCl concentrations on root length of different varieties of *Sorghum* seedlings.

For each *Sorghum* NaCl treatment, values within a column sharing same letter comparing NaCl treatments are not significantly different at p<0.05 (Fishers LSD). Each value represented as mean ± SE are the mean of three replications.

### Effects of salt stress on relative water content

In relation to relative water content, the results point to a decreasing trend upon salinity stress, with the decrease depending on the level of NaCl stress applied (Figure 3). As expected, in the absence of NaCl stress, the relative water content increased over time in all studied *Sorghum* varieties. Plants grown under moderate water stress treatment of 100 mM NaCl demonstrated the highest diversity relative water content values. In contrast, the three varieties irrigated with 300 mM NaCl had a significantly reduced relative water content, compared to the control plants. At this concentration, very small shoots formed all varieties Sc Sila and Serena, resulting in the lowest water content.

### Effects of salt stress on total chlorophyll content

Results from (Figure 4) shows an inverse relationship between NaCl induced drought stress responses and total chlorophyll content values for all *Sorghum* varieties tested. Differences for chlorophyll content values were also observed among varieties. At the beginning of the experiment, total chlorophyll content across the varieties was similar ranging from 9.37 to 9.44 mg/g FW. Imposition of moderate salinity stress conditions of 100 mM NaCl caused a slight decrease of chlorophyll content which ranged from 5.8730 mg/g FW for Serena to 8.15 mg/g FW for Gadam. In severe salt stress conditions of 100 mM NaCl, significant decrease of total chlorophyll content was also observed among the three varieties with Gadam having the highest (4.54 mg/g FW), while Serena had the least (3.13 mg/g FW). Among the varieties exposed to severe salt stress, varieties Gadam retained relatively high chlorophyll content when compared with Sc Sila and Serena.

**Figure 4:**
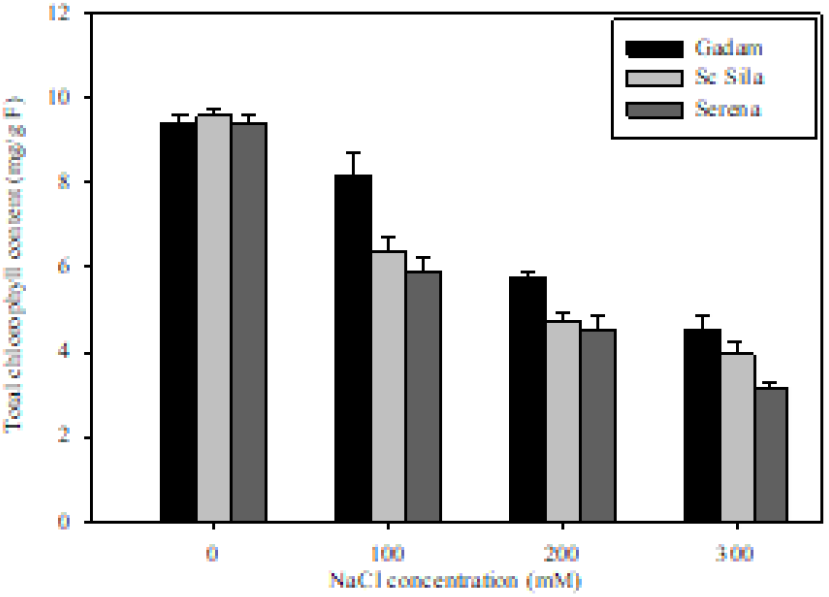
Effects of salt stress on total chlorophyll content.

## DISCUSSION

Salinity stress is one of the most serious adverse factors that affect growth and productivity of salt-sensitive plant species, including *Sorghum* [9,10]. Increased irrigation and the predicted climate change are expected to increase the severity and frequency of soil salinization, especially in arid and semi-arid areas where cultivation of *Sorghum* occurs. Developing varieties with improved salinity tolerance will therefore help in sustaining increased productivity under agricultural areas that are prone to salinity stress. The challenging aim of breeding for enhanced yield under salinity stress is certainly dependent on existing selection methodologies and robust screening of salt-tolerant *Sorghum* germplasm. Due to complex inheritance of traits coupled with wide variations in environmental conditions, the applied screening methods are time-consuming and inefficient. Hence, it is essential to define selection criteria to be employed for identification of tolerant varieties. Because of this, seed germination potential under salt stress has demonstrated to be a reliable tool for estimation of genetic tolerance. This strategy allows the selection of more salt tolerant plant varieties at early development phases, therefore allowing for greater cost-effectiveness and time-efficient of all subsequent breeding activities. This research aimed to determine the potential for seed germination of three *Sorghum* varieties subjected to different concentrations of salinity stress.

Our findings reveal that salinity stress influences significantly all *Sorghum* characteristics associated with germination and early seedling development, with salinity tolerance depending on the stress level of the selected varieties. Nevertheless, the varieties responded differently to the varying concentrations of salinity. As regards germination, these variations were linked to the proportion of final germination as well as the rate of germination. Our findings are comparable with those of other research which show that salinity stress acts as an inhibitor by stopping germination, without loss of viability at high stress levels, and by delaying germination at concentrations where the process is not completely avoided [11,12]. Salt stress affects seed germination mainly by reducing the soil solution’s osmotic potential to delay seed water absorption, causing embryo toxicity to sodium and/or chloride, or changing protein synthesis. Again, poor seedling emergence might be caused by hypocotyls mortality associated with the salt accumulation at the soil surface [13]. Even though germination rate of all varieties was considerably altered by high level of salinity stress, the most severe effects were recorded in genotype Serena and Sc Sila, whereas Gadam had the best performance with regard to this trait. Our results also revealed that increasing salinity stress concentrations progressively reduced development of seedling, expressed as decreased lengths of root and that the salt effect depended on salinity level and variety. As anticipated, these length decreases were the greatest at the highest salinity levels and most probably mirror the toxic effects combined with insufficient water uptake and nutrients [14,15]. Furthermore, there was significant variation in reduced length between organs and between varieties. The shoot and root lengths were differentially affected by salinity stress, with roots being more seriously affected even under low NaCl concentrations. The observed effects of salinity stress on roots could be attributed to the to the direct exposure salinity. We hypothesized that Gadam could perform satisfactorily in soils with less severe salinity.

An essential strategy for plant tolerance to NaCl stress is the ability to retain high water status during salinity stress. Decrease in relative water content as in response to salinity stress has been reported in a wide variety of plants [16,17]. This variation in the capacity of plants to hold water can be ascribed to their differential capacity to absorb water from the soil by creating a reduced potential gradient of water from the soil and also because of the distinction in the capacity of the plant varieties to adjust and maintain osmotic turgor in the tissues. The differences in relative water content in all varieties observed in our study could be correlated with their different ability of water absorption from soil. The decline in relative water content recorded was a main factor that caused decreased growth responding to osmotic stress in the *Sorghum* plants. Under salt stress, Sc Sila and Serena varieties were more affected by the decrease in relative water content than Gadam. This suggested that the three *Sorghum* varieties had different sensitivity when subjected NaCl induced salt stress. The increased water retention capability found in the Gadam could play a crucial part in crop’s survival under salinity stress.

Chlorophyll content strongly depends on the species’physiological responses and their ability to tolerate stress [18]. Measurement of chlorophyll content is one of the most effective indicators for salinity tolerance identification of *Sorghum* [19]. In our study, salinity stress caused an increase in total chlorophyll content in the leaves of all *Sorghum* varieties although the decrease was differed in terms of variety and the stress level. These observations explained why the total chlorophyll content of all decreased under salinity stress. The decrease in total chlorophyll content could be due the accumulation of Na^+^ and Cl^−^ ions which hinders the process of chlorophyll synthesis by influencing the activity of some enzymes containing Fe^3+^. Besides this, the decrease in chlorophyll contents might be related to an increase of chlorophyll degradation or a decrease of chlorophyll synthesis [20].

## CONCLUSION

In conclusion, our study findings confirm that *Sorghum’s* tolerance to salinity stress is highly variety-dependent, as manifested by differentiation in terms of germination and early seedling development under salinity stress. In addition, our findings point to the existence of large genetic variation for this specific trait among the different *Sorghum* varieties. Given the need to identify new sources of salt tolerance, the observed varietal variation indicates that a larger number of *Sorghum* varieties should be considered in future research. Our findings underline that variability in the stress response may be readily explored during the germination phase and early seeding development, for reliable selection of salt-tolerant genotypes at early growth stages. Finally, the findings indicate Gadam’s supremacy in tolerating salinity stress, thus suggesting the possibility for its cultivation under salt stress conditions as well as its suitability for use as germplasm material in future *Sorghum* breeding programmes.

## ACKNOWLEDGMENTS

The authors thank Pwani University for providing materials, reagents and lab space to conduct this study.

## REFERENCES

1. Vaidyanathan H, Sivakumar P, Chakrabarty R, Thomas G. Scavenging of reactive oxygen species in NaCl-stressed rice (Oryza sativa L.)-differential response in salt-tolerant and sensitive varieties. Plant Sci. 2003;165:1411–1418.

2. Acosta-Motos J, Ortuño M, Bernal-Vicente A, Diaz-Vivancos P, Sanchez-Blanco M. Plant responses to salt stress: Adaptive mechanisms. agronomy. 2017;7:18.

3. HanumanthaRao B, Nair RM, Nayyar H. Salinity and high temperature tolerance in mungbean [Vigna radiata (L.) Wilczek] from a physiological perspective. Front Plant Sci. 2016;7:957.

4. Debez A, Belghith I, Pich A, Taamalli W, Abdelly C. High salinity impacts germination of the halophyte cakile maritima but primes seeds for rapid germination upon stress release. Physiol Plant. 2018;164:134–144.

5. Bibi A, Sadaqat HA, Tahir MHN, Akram HM. Screening of *Sorghum* (*Sorghum* bicolor Var Moench) for drought tolerance at seedling stage in polyethylene glycol. J Anim Plant Sci. 2012;22:671–678.

6. Ramirez-Villegas J, Jarvis A, Läderach P. Empirical approaches for assessing impacts of climate change on agriculture: The EcoCrop model and a case study with grain *Sorghum*. Agric For Meteorol. 2013;170:67–78.

7. Calviño M, Messing J. Sweet *Sorghum* as a model system for bioenergy crops. Curr Opin Biotechnol. 2012;23:323–329.

8. Arnon DI. Copper enzymes in isolated chloroplasts. Polyphenoloxidase in beta vulgaris. Plant Physiol. 1949;24:1–15.

9. Netondo GW, Onyango JC, Beck E. *Sorghum* and salinity. Crop Sci. 2004;44:797–805.

10. De La Rosa-Ibarra M, Maiti RK. Biochemical mechanism in glossy *Sorghum* lines for resistance to salinity stress. J Plant Physiol. 1995;146:515–519.

11. Khayamim S, Rouzbeh F, Poustini K, Sadeghian SY, Abbasi Z. Seed germination, plant establishment, and yield of sugar beet genotypes under salinity stress. J Agr Sci Tech. 2018;16:779–790.

12. Bajji M, Kinet JM, Lutts S. Osmotic and ionic effects of NaCl on germination, early seedling growth, and ion content of atriplex halimus (Chenopodiaceae). Can J Bot. 2002;80:297–304.

13. Dias LS, Dias AS, Pereira IP. Ionic effects of NaCl counter osmotic inhibition of germination and seedling growth of Scorzonera hispanica and subsequent plantlet growth is not affected by salt. Botany. 2015;93:485–496.

14. Ouji A, El-Bok S, Mouelhi M, Younes MB, Kharrat M. Effect of salinity stress on germination of five Tunisian lentil (Lens culinaris L.) genotypes. ESJ. 2015;11: 21.

15. Majid A, Mohsen S, Mandana A, Saeid JH, Ezatollah E. The effects of different levels of salinity and indole-3-acetic acid (IAA) on early growth and germination of wheat seedling. J Stress Physiol Biochem. 2013;9:329–338.

16. Nxele X, Klein A, Ndimba BK. Drought and salinity stress alters ROS accumulation, water retention, and osmolyte content in *Sorghum* plants. S Afr J Bot. 2017;108:261–266.

17. Silva ED, Ribeiro RV, Ferreira-Silva SL, Viégas RA, Silveira JAG. Comparative effects of salinity and water stress on photosynthesis, water relations and growth of Jatropha curcas plants. J Arid Environ. 2010;74:1130–1137.

18. Chaves MM, Flexas J, Pinheiro C. Photosynthesis under drought and salt stress: regulation mechanisms from whole plant to cell. Ann Bot. 2009;103:551–560.

19. Romero L, Belakbir A, Ragala L, Ruiz JM. Response of plant yield and leaf pigments to saline conditions: Effectiveness of different rootstocks in melon plants (Cucumis melo L.). J Soil Sci Plant Nutr. 1997;43:855–862.

20. Netondo GW, Onyango JC, Beck E. *Sorghum* and salinity. Crop Sci. 2004;44:797–805.

